# Hybrid Gated Fusion: A Multimodal Deep Learning Framework for Protein Function Annotation

**DOI:** 10.64898/2026.04.14.718564

**Authors:** Zijian Zhou, Daniel WA Buchan

**Affiliations:** Department of Computer Science, University College London, London, UK

**Keywords:** protein function prediction, multimodal learning, deep learning, bilinear fusion, Gene Ontology

## Abstract

Protein function annotation requires integrating diverse biological signals, yet existing multimodal methods often struggle with missing inputs and redundant information. We present Hybrid Gated Fusion, a multimodal architecture that combines intrinsic protein features, including sequence and structure, with extrinsic functional context from text and interaction networks. Rather than weighting all modalities equally, the model uses bilinear gating to assess both the informativeness of each modality and its agreement with the others, while auxiliary supervision reduces modality dominance and preserves useful signal in weaker modalities. On the CAFA3 benchmark, a single Hybrid Gated Fusion model achieves state-of-the-art performance in Biological Process (*F*_max_ = 0.601) and Cellular Component (*F*_max_ = 0.706), while remaining competitive in Molecular Function (*F*_max_ = 0.702). Analysis of the learned gates shows that interaction networks and text often provide complementary functional signals, whereas structural features are down-weighted when redundant but remain valuable under sparse-input settings. These results establish Hybrid Gated Fusion as a robust and scalable framework for genome-scale protein function annotation.

**Availability and implementation:** Source code and reproduction scripts are freely available at https://github.com/psipred/PFP. Pre-computed embeddings, data splits, and model checkpoints are deposited at https://doi.org/10.5281/zenodo.19498341.

## 1. Introduction

Assigning biological function to proteins is essential for interpreting genomes, reconstructing cellular pathways, and identifying therapeutic targets, yet the gap between known sequences and experimentally characterised functions continues to widen. High-throughput sequencing has expanded the catalogue of protein sequences to approximately 246 million entries in UniProt (Consortium, 2025), but only a small fraction carry experimentally validated functional annotations, creating a pressing need for accurate and scalable computational predictors. The most widely adopted formalism for describing protein function is the Gene Ontology (GO) (Ashburner et al., 2000), which organises functional descriptors into three structured hierarchical ontologies, Biological Process (BPO), Molecular Function (MFO), and Cellular Component (CCO), each posing distinct prediction challenges due to differences in label granularity and annotation density.

Protein function arises through both intrinsic determinants and extrinsic context, and these signals can be encoded through different deep learning approaches. Large protein language models (PLMs), including ProtTrans (Elnaggar et al., 2022) and ESM (Rives et al., 2021), capture evolutionary and functional semantics from sequence; structure-informed encoders leveraging AlphaFold predictions (Jumper et al., 2021) encode geometric constraints; and network-based approaches exploiting protein– protein interaction (PPI) graphs (You et al., 2021) provide relational evidence complementary to sequence and structure. These complementary signals motivate *multimodal learning* approaches to GO annotation (Kulmanov et al., 2018; Boadu et al., 2023) and are also reflected in emerging foundation models that jointly model sequence, structure, and function (Hayes et al., 2025).

Despite this progress, practical multimodal prediction faces two persistent limitations. First, many frameworks assume complete modality availability at inference time, yet real-world coverage is uneven: protein sequence data is universally available, whereas highquality structures, curated text, and verified interaction networks are frequently missing. Remedies such as zero-filling, imputation, or discarding incomplete samples can inject noise, bias fusion behaviour, or reduce the effective training data (Ma et al., 2022). Second, fusion mechanisms often trade off expressiveness against data efficiency: simplistic aggregation may fail to exploit cross-modal complementarities, whereas more expressive fusion architectures can be harder to train robustly and may be more prone to overfitting in small-sample settings (Baltrusaitis et al., 2019; Stahlschmidt et al., 2022). These constraints necessitate deep learning approaches with strong inductive bias and explicit robustness to missing or unreliable evidence.

To evaluate GO-based protein function prediction under realistic generalisation, the Critical Assessment of Functional Annotation (CAFA) challenge has become the standard benchmark for temporal generalisation (Zhou et al., 2019). In this setting, test proteins lack experimental annotations at the prediction cutoff and are evaluated only against annotations added afterward. Because these proteins are often less completely characterised than the training set, CAFA provides a natural benchmark for methods that must remain robust under incomplete evidence.

In this work, we present Hybrid Gated Fusion, a multimodal framework for GO annotation that integrates intrinsic protein evidence (sequence and structure) with extrinsic functional context (text and interaction networks) under incomplete availability of all input features. The model first applies a learnable Bilinear Gated Early Fusion module to estimate how informative each available modality is and how well it agrees with the others, producing a fused representation that emphasizes complementary evidence. It then uses Residual Late Fusion, in which modality-specific auxiliary predictions are combined with the same gating weights so that decision-level contributions remain aligned with feature-level evidence quality. This design supports dynamic masking for missing inputs, reduces spurious confidence when evidence is noisy or absent, and achieves state-of-the-art performance on CAFA3, particularly for BPO and CCO, without relying on computationally expensive Multiple Sequence Alignment (MSA) generation. In the results we also analyse the learned fusion behaviour, showing how modality weights reflect marginal utility across evidence sources and how auxiliary supervision mitigates modality dominance to preserve discriminative capacity for sparse inputs.

## 2. Materials and Methods

We introduce a multimodal architecture for protein function prediction designed to be (i) resilient to missing feature modalities, (ii) parameter-efficient, and (iii) interpretable through modality-specific gating. We formulate the task as a multi-label prediction problem over heterogeneous inputs (see Section 2.1) and evaluate our approach on the standard CAFA3 benchmark using temporally split train/validation/test sets (Sec. 2.2). Figure 1 summarizes the pipeline in five stages. First, four complementary evidence sources, protein sequence, functional text, protein structure, and proteinprotein interactions, are encoded into a shared latent space (Sec. 2.3). Second, the resulting embeddings are standardized and paired with a modality-availability mask so that the model can operate on arbitrary evidence subsets without imputation (Sec. 2.4). Third, the **Bilinear Gated Early Fusion** module estimates modality reliability and cross-modal agreement to construct a fused latent representation (Sec. 2.5). Fourth, modality-specific auxiliary heads and **Residual Late Fusion** preserve discriminative signal in each modality while reusing the early-fusion weights for consistent late-stage aggregation (Sec. 2.6). Finally, the early and late pathways are combined to produce the final multi-label GO prediction (Sec. 2.6).

**Figure 1.**
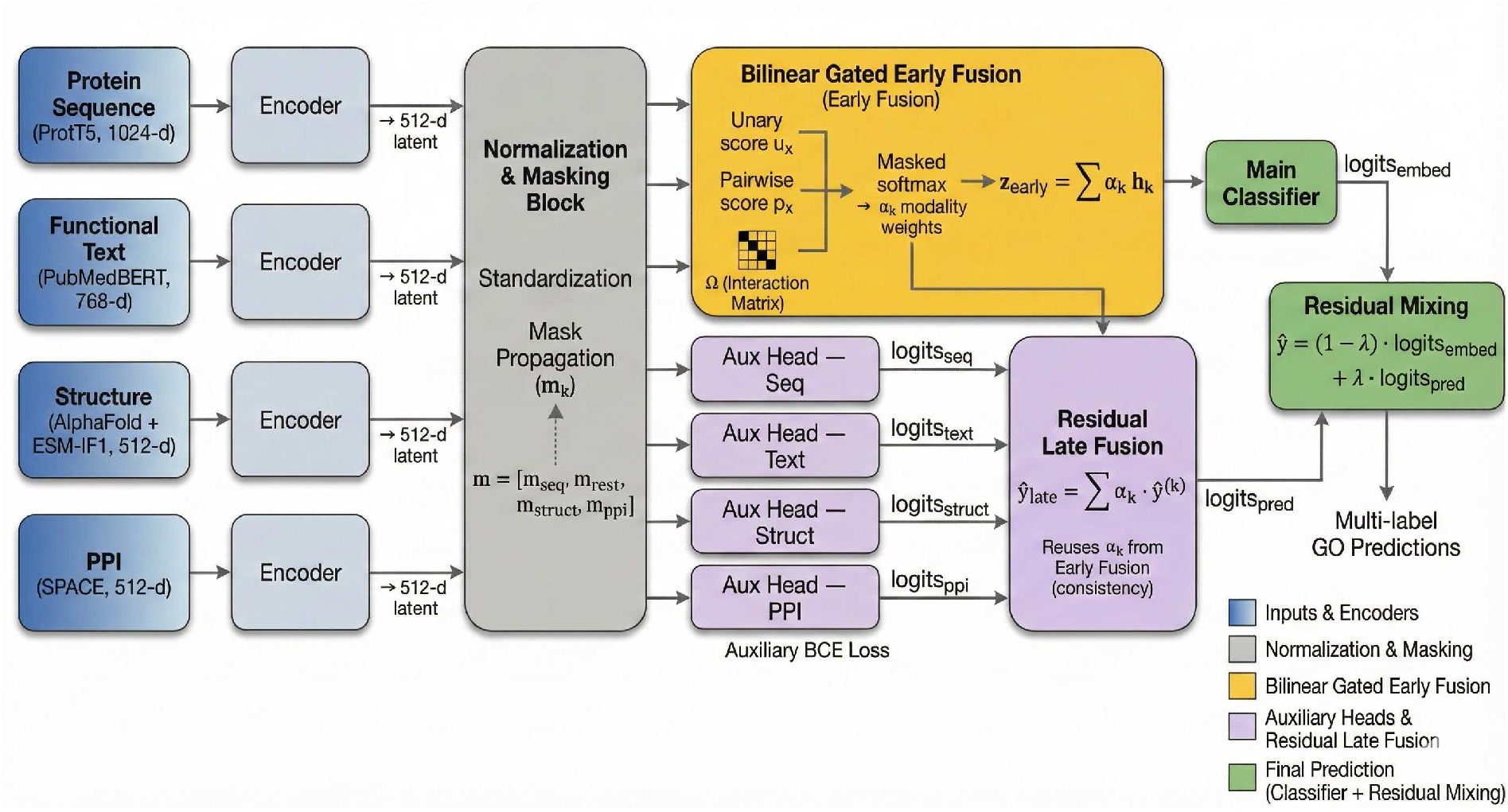
Hybrid Gated Fusion architecture and prediction pipeline. The pipeline contains five stages that correspond to the Method subsections. Inputs and Encoders (Sec. 2.3): sequence, text, structure, and PPI evidence are encoded with ProtT5, PubMedBERT, ESM-IF1, and SPACE, respectively, then projected into a shared latent space. **Normalization and Masking** (Sec. 2.4): modality embeddings are standardized, and missing inputs are handled with a binary availability mask m and mask propagation. **Bilinear Gated Early Fusion** (Sec. 2.5): unary modality scores and pairwise interactions estimate modality relevance and cross-modal agreement, yielding the fused representation z_early_. **Auxiliary Heads and Residual Late Fusion** (Sec. 2.6): modality-specific auxiliary heads produce logits that are aggregated with the same attention weights *α*_*k*_, providing consistency-aware late fusion. **Final Prediction** (Sec. 2.6): the main classifier and residual mixing module combine early- and late-fusion outputs to generate multi-label GO predictions.

### 2.1. Problem Formulation

We formulate protein function prediction as a multi-label classification problem over a heterogeneous set of biological feature modalities. Let 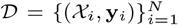 denote a dataset of *N* proteins, where y_*i*_ ∈ {0, 1}^*C*^ represents binary annotations for *C* possible Gene Ontology (GO) terms. Each protein may be associated with any subset of the four modalities ℳ_all_ = {Seq, Text, Struct, PPI}, and its available inputs are

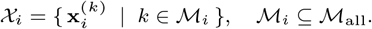

Due to incomplete experimental coverage, modality availability varies across proteins. We therefore introduce a binary mask 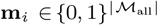 that indicates which modalities are present. The learning objective is to estimate a function

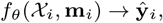

that maps arbitrary subsets of modalities to GO predictions in a manner that is robust to missing inputs and scalable across thousands of GO labels.

### 2.2. Benchmark Dataset

We evaluate our approach on CAFA3 using the processed temporal splits provided by Oliveira et al. (2023). These splits define the train, validation, and test partitions used throughout the paper and preserve the time-delayed evaluation setting of CAFA.

The CAFA3 benchmark poses significant challenges for multimodal learning: (i) extreme label sparsity, (ii) high cardinality (thousands of candidate terms), and (iii) the “open-world” nature of protein function, where absence of an annotation does not imply a negative. Additionally, while sequence data is available for all targets, structural and interaction data is not always available for all input proteins. A detailed breakdown of the data distribution is provided in Appendix A (Table 3).

### 2.3. Feature Extraction Protocols

We represent protein function prediction using complementary evidence from intrinsic attributes (sequence and structure) and extrinsic context (textual metadata and interaction networks). Each modality is encoded into a fixed-dimensional embedding using a pretrained encoder, and these embeddings form the inputs to our fusion model. Full preprocessing and implementation details are provided in Appendix B.

Protein sequences were encoded with ProtT5 (Elnaggar et al., 2022), a large protein language model trained on amino-acid sequences. Its embeddings capture statistical patterns in sequence data that reflect underlying evolutionary constraints and biochemical properties, making sequence a strong baseline signal that is available for all proteins.

Protein structure is encoded from AlphaFold-predicted coordinates (Jumper et al., 2021) using ESM-IF1 (Hsu et al., 2022), an inversefolding encoder based on Geometric Vector Perceptrons (Jing et al., 2021). We use this encoder in a sequence-agnostic manner by providing backbone geometry only, so that the structural modality reflects three-dimensional organisation, such as spatial arrangement and geometric context, rather than re-encoding sequence-derived cues already available through the sequence model.

Textual features are derived from curated UniProtKB metadata (Consortium, 2018) and encoded with PubMedBERT (Gu et al., 2021). Because UniProt records are continuously enriched, using current entries for proteins in a temporally split test set risks data leakage. To prevent this, we retrieve historical UniProt entries from the UniSave archive (Leinonen et al., 2006) for test set proteins, retaining only records that predate the CAFA3 training cutoff; train and validation proteins use current records. Structured fields are extracted, sanitised, and concatenated into a single input string (Appendix B). The resulting embedding provides a semantic representation that aligns natural-language functional descriptions with GO terminology.

Interaction context is captured from the STRING protein– protein interaction network database (Szklarczyk et al., 2023) using SPACE embeddings (Hu et al., 2025). These embeddings summarise global network topology and neighbourhood structure, providing complementary context for pathway- and system-level functions that are not readily inferred from an isolated protein sequence or structure.

### 2.4. Normalization and Dynamic Missing Modality Handling

To mitigate disparities between the size of the feature vectors arising from different pre-trained encoders, embeddings are mapped to a shared dimension *d*_*model*_ via shallow projection blocks *f*_*k*_(·), each consisting of a single Linear layer followed by Layer Normalization and a ReLU activation. This processing ensures that subsequent fusion layers operate on consistent dimension vectors, which helps stabilise gradients while preserving modality-specific biological variance.

We address missing modality availability through a set-theoretic masking approach. We define a binary availability mask vector 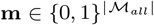, such that *m*_*k*_ = 1 if *k* ∈ ℳ_*i*_ and 0 otherwise. Rather than imputing missing features, we employ zero-padding combined with strict mask propagation. The mask m is utilised to (1) block gradient updates to encoders of missing modalities and (2) ensure missing modalities contribute exactly zero to the attention scores in the subsequent fusion layers.

### 2.5. Bilinear Gated Early Fusion

The early-fusion module assigns each available modality a weight based on two signals: its standalone informativeness and its compatibility with the other available modalities. In protein function prediction, modalities often exhibit complex dependencies. Ideally, the model should distinguish between *redundant* signals (e.g., functional text merely explicitly stating motifs present in the sequence) and *interactive* signals (e.g., 3D structural information revealing enzymatic pockets not obvious from primary sequence conservation alone).

The fusion mechanism computes a scalar attention weight *α*_*k*_ for each modality *k* via a biologically grounded hybrid scoring function: First, we compute a single modality score *u*_*k*_, estimating the intrinsic informational quality of modality *k* in isolation. This allows the model to down-weight noisy experimental data (e.g., high-throughput PPIs) in favor of high-confidence signals (e.g., evolutionary sequence profiles):

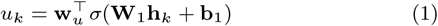

where *σ* denotes the GELU activation function. Simultaneously, we model second-order interactions to capture cross-modal biological context. We project features into a subspace via W_*P*_ and introduce a learnable Interaction Matrix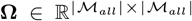. The pairwise contribution *p*_*k*_ is calculated as:

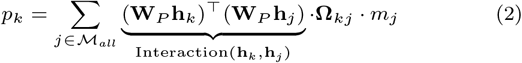

Here, Ω_*kj*_ acts as a learnable coefficient modulating the interaction between modalities *k* and *j*. By scaling feature compatibility, this term enables the network to learn data-driven aggregation strategies, promoting complementary signals while dampening redundant ones. The critical output of this mechanism is the normalised attention weight *α*_*k*_, which serves as the model’s definitive measure of biological relevance for each data source. This weight is derived from an intermediate score *s*_*k*_ that synthesises the standalone quality *u*_*k*_ and contextual support *p*_*k*_ via a learnable balance scalar *γ*:

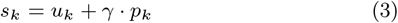

To ensure a valid probability distribution over dynamic subsets of biological evidence, we apply a masked softmax to these scores:

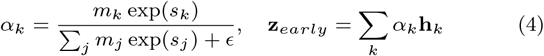

Crucially, *α*_*k*_ serves two purposes: it constructs the fused representation z_*early*_ and acts as a reliability gate for the downstream auxiliary predictions. This keeps the late-fusion ensemble aligned with the feature relevance estimated at the early-fusion stage.

### 2.6. Auxiliary Heads and Residual Late Fusion

In protein function prediction, sequence-derived features are available for nearly all proteins and during training this data modality can come to dominate optimization, even when structure, text, or PPI evidence provides complementary functional information. This creates a modality-dominance problem: sparse modalities may be under-utilised during training despite carrying biologically useful signals. To address this, we equip each input modality track with an auxiliary prediction head and introduce a Residual Late Fusion pathway. Together with Bilinear Gated Early Fusion, these components form the overall Hybrid Gated Fusion architecture.

Each modality encoder is equipped with a linear classifier that produces logits ŷ^(*k*)^ ∈ ℝ^*C*^. These auxiliary heads are trained jointly with the main model so that each modality embedding remains independently predictive of protein function. This additional supervision helps preserve discriminative information in sparse modalities, such as structure and PPI, rather than allowing them to be treated as weak secondary signals during optimization.

Residual Late Fusion then links feature-level weighting to prediction-level aggregation. We reuse the modality attention weights *α*_*k*_ derived from the Bilinear Gated Early Fusion module to combine the auxiliary predictions, so that modalities judged more informative during feature fusion also contribute more strongly at the decision level. The late-fusion ensemble ŷ_*late*_ is computed as:

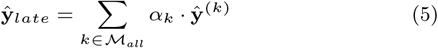

This consistency-aware aggregation keeps the late-fusion pathway aligned with the evidence weighting learned upstream, allowing the model to amplify informative modalities while suppressing noisy or weakly supported ones.

The final model prediction ŷ combines the early-fusion classifier, which operates on the joint latent representation z_*early*_, with the late-fusion ensemble through a learnable residual connection. Let *λ* = *σ*(*β*) ∈ (0, 1) be a gating coefficient derived from a learnable scalar *β*. The final output is defined as:

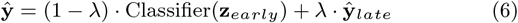

This formulation allows the network to dynamically hedge its predictions: it can rely on complex, non-linear biological mechanisms captured in the latent space (z_*early*_) while maintaining the robustness of an ensemble of independent evidence sources (ŷ_*late*_).

### 2.7. Optimization

The model is trained using a joint Binary Cross Entropy (BCE) objective. To prevent feature collapse in the auxiliary branches, we introduce a learnable parameter *η* to weight the auxiliary supervision:

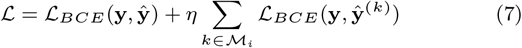

### 2.8. Evaluation Metrics

We evaluate the performance using the standard CAFA metrics: Maximum F-measure (*F*_max_) and Semantic Distance (*S*_min_), computed via the official CAFA-evaluator toolkit (Piovesan et al., 2024). We prioritize the weighted metrics (*wF*_max_, *wS*_min_) as the more rigorous evaluation standard. By scaling performance based on Information Accretion (IA), these metrics prevent models from inflating scores through the prediction of broad, shallow terms (e.g., “cellular process”). *wF*_max_ rewards the retrieval of information-rich terms, while *wS*_min_ quantifies the information error by combining Remaining Uncertainty (false negatives) and Misinformation (false positives). To facilitate straightforward benchmarking against prior state-of-the-art methods, we also report the unweighted counterparts (*F*_max_, *S*_min_). Since official IA weights were not provided with the CAFA3 dataset, we computed them directly from training annotations following standard definitions. Complete mathematical definitions for all metrics and weight calculations are provided in the Appendix C.

### 2.9. Experimental Setup

We implemented the framework in PyTorch and trained all models on a single NVIDIA RTX 4070 Ti Super (16GB), fixing the random seed to 42 for reproducibility. Optimization used AdamW (learning rate 1 × 10^−3^, weight decay 1 × 10^−2^) with a 10% linear warmup followed by cosine annealing. Training ran with a batch size of 32 for up to 50 epochs, utilising early stopping (patience 5) and modality-specific dropout (0.1) to enhance robustness. Input dimensions were standardized prior to fusion (*d*_*model*_ = 512): sequence (1024-d), text (768-d), and structure/PPI (512-d).

## 3. Results

### 3.1. Contribution of the hybrid fusion architecture

In order to assess the robustness of our framework under realworld biological constraints, where evidence is often incomplete, we evaluated the specific contribution of the residual late-fusion pathway. We compared the full **Hybrid Gated Fusion** model against an ablated Bilinear Gated baseline (early fusion only) by dynamically masking input channels at inference time. This experiment targets *modality dominance*: without an explicit mechanism to preserve non-sequence channels, training can over-optimise for sequence while leaving sparse modalities (e.g., Structure and PPI) weakly discriminative.

Figure 2A–C summarises the resulting *wF*_max_ trends across each GO domain. In the absence of sequence, the early-fusion baseline degrades sharply, most notably for structure-only and PPI-only inference, indicating that these channels contribute limited signal when the dominant sequence cue is removed. By contrast, the hybrid model shows substantially improved resilience in these sparse regimes. For example, in BPO with structure-only input, *wF*_max_ increases from 0.256 to 0.424 (+65%), and *wS*_min_ improves from 22.92 to 17.85, indicating not only higher accuracy but also predictions that are semantically closer to the ground truth. Similar recovery is observed for PPI-only settings, consistent with the auxiliary supervision objectives preventing these encoders from collapsing into non-informative representations.

**Figure 2.**
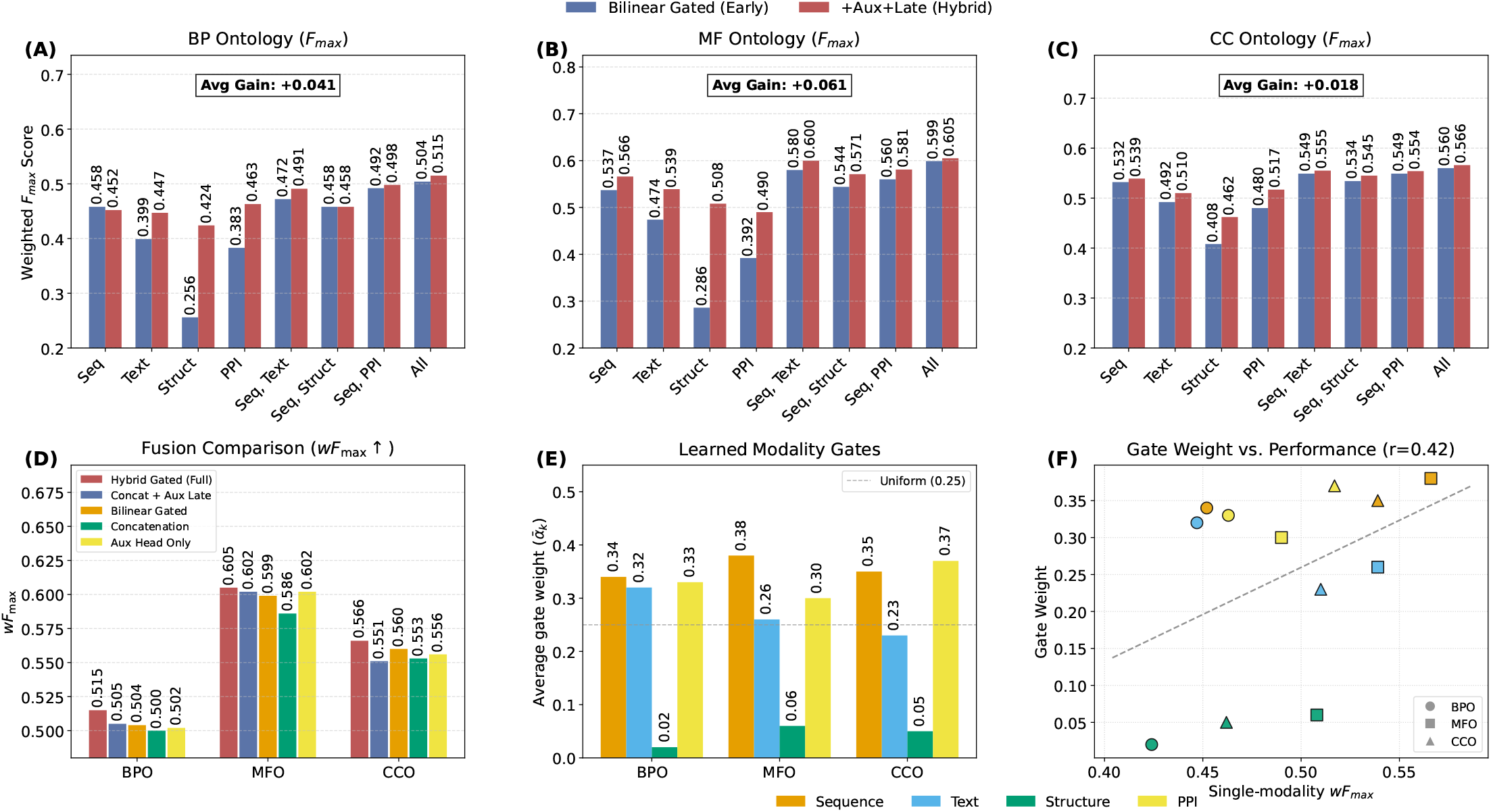
Modality robustness, fusion ablation, and learned gating dynamics. **(A–C)** Modality-subset robustness under missing evidence: *wF*_max_ for the *Bilinear Gated* baseline and the proposed *Hybrid Gated Fusion* model across **BPO** (A), **MFO** (B), and **CCO** (C). The x-axis denotes the subset of input modalities provided at inference using the paper shorthand `Seq`, `Text`, `Struct`, `PPI`, and `All` (e.g., `Seq` for sequence-only inference). `Avg Gain` reports the mean improvement of the hybrid model over the baseline across all modality combinations within each ontology. **(D)** Fusion ablation on weighted metrics: comparison of fusion configurations using *wF*_max_ (higher is better). **(E–F)** Analysis of learned gating dynamics. **(E)** Global average gating coefficients 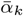 computed over the CAFA3 test set using the full-modality configuration (ℳ_all_); the dotted horizontal line indicates uniform weighting (0.25). **(F)** Relationship between learned gate weights and single-modality predictive performance (*wF*_max_): each point corresponds to a modality–ontology pair, colour-coded by modality (matching panel E) and shape-coded by ontology (circle: BPO; square: MFO; triangle: CCO). The dashed diagonal line shows the linear regression trend (*r* = 0.42).

In full-modality inference (M_*all*_), the hybrid model remains competitive and yields modest gains in BPO (+2.2%), MFO (+1.0%), and CCO (+1.1%), suggesting that the residual pathway acts as a stabilising ensemble component rather than trading off peak performance for robustness. Sequence-only performance is largely preserved, with only minor differences between the two models, whereas the baseline exhibits pronounced drops across other single-modality configurations. Collectively, these results are consistent with a modality-dominance effect: without auxiliary supervision, early fusion permits the model to satisfy the training objective predominantly through the sequence channel, leaving sparse encoders under-trained. The hybrid architecture mitigates this by enforcing independent discriminability for each modality, thereby preserving predictive capacity when dominant inputs are unavailable. Full weighted results for all masked-modality configurations (including *wS*_min_) are reported in Supplementary Appendix G, with unweighted counterparts in Appendix H.

#### 3.1.1. Analysis of Fusion Mechanisms

To quantify the contribution of the fusion design, we perform a controlled ablation study targeting (i) *bilinear interactions* in the early fusion gate and (ii) *consistency-aware aggregation* for coordinating early and late pathways. We compare the full Hybrid Gated model against four variants: *Bilinear Gated* (early pathway only), *Concatenation* (dense projection replacing bilinear interactions), *Aux Head Only* (late pathway only with learnable gating), and *Concat + Aux Head Only* (hybrid residual baseline without shared weights). All variants use identical modality encoders, prediction heads, and training hyperparameters. Weighted results are summarised in Figure 2D; full weighted and unweighted tables are reported in Appendix E and Appendix F, respectively.

Overall, the full **Hybrid Gated** model achieves the strongest weighted performance across all three ontologies, reaching *wF*_max_*/wS*_min_ of **0.515/16.08** (BPO), **0.605/5.98** (MFO), and **0.566/6.22** (CCO) (Appendix E). Comparing early-fusion designs, *Bilinear Gated* consistently outperforms *Concatenation*, indicating that the gains arise from explicit pairwise interactions rather than increased fusion capacity (details in Appendix D).

When evaluated in isolation, *Bilinear Gated* (early-only) and *Aux Head Only* (late-only) show comparable performance, suggesting that both pathways capture useful and complementary functional signal. However, hybridising the pathways without coordination (*Concat + Aux Head Only*) does not reliably improve over its constituents. In contrast, the full model consistently improves both accuracy and semantic quality, supporting the role of *consistency-aware aggregation*: reusing early-stage weights *α*_*k*_ aligns late-stage contributions with feature compatibility, allowing the model to suppress noisy auxiliary predictions when the early gate indicates low information quality.

#### 3.1.2. Interpretability: Learned gating dynamics

To examine how the model allocates weight across biological evidence sources, we analyse the softmax-normalised gating coefficients *α*_*k*_ from the fusion layer (Eq. 4). Specifically, we compute the global mean weight 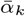 over the CAFA3 test set under full-modality inference (Fig. 2E–F). Appendix J complements this cohort-level analysis with protein-level case studies of strongly polarized gating.

The learned gates broadly follow standalone modality utility (*r* = 0.42; Fig. 2F), but they also reflect *marginal* rather than absolute value. Sequence receives the largest weight in BPO and MFO, for example reaching 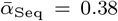 in MFO, consistent with sequence being the strongest general-purpose signal. However, the weighting pattern also highlights complementary evidence: in CCO, PPI is assigned slightly more weight than sequence (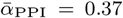 vs. 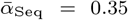), consistent with interaction data contributing localisation-specific context that is not fully captured by sequence alone. Text occupies an intermediate role across all three ontologies 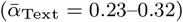, suggesting that curated functional descriptions provide broadly useful but not dominant evidence. By contrast, structure is consistently down-weighted 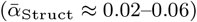 despite competitive single-modality MFO performance, indicating that under full-modality inference it contributes less incremental information once sequence, text, and PPI are already available.

### 3.2. Modality Contribution Analysis

This analysis quantifies the marginal value of adding each auxiliary modality on top of the universally available sequence signal. We anchor the comparison on sequence because ProtT5 embeddings are available for all CAFA3 targets, whereas text, structure, and PPI inputs have incomplete coverage (Table 3); this makes sequence the fairest deployable reference rather than an arbitrary baseline. Accordingly, the *Sequence Only* model uses the same ProtT5 encoder and classifier architecture as the multimodal system but removes auxiliary modalities and fusion, so gains in the bi-modal rows isolate the contribution of adding text, structure, or PPI. Unlike the inference-time masking used in the ablation study (Section 3.1), each subset model is retrained for its available inputs, providing an upper bound for that evidence configuration. The results, summarised in Table 1, reveal three distinct biological patterns. Unweighted metrics are detailed in Appendix I.

**Table 1.**
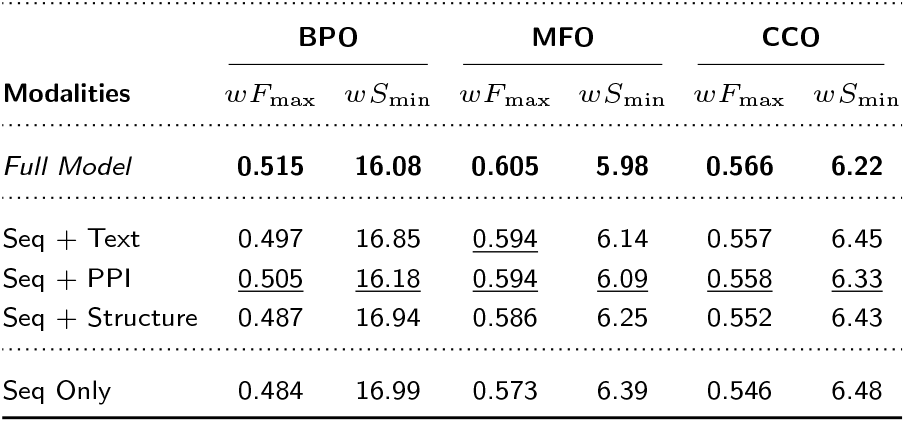
Modality Contribution Analysis. Models were retrained on specific modality subsets to estimate the marginal value of each auxiliary modality when added to the universally available sequence signal. Bimodal rows therefore report gains over the *Sequence Only* anchor, while the **Full Model** tests whether combining all modalities yields further benefit. **Metrics**: *wF*_max_ (↑) and *wS*_min_ (↓). Best results are bold; second-best are <u>underlined</u>.

| Modalities | BPO |  | MFO |  | CCO |  |
| --- | --- | --- | --- | --- | --- | --- |
| | $wF_{\max}$ | $wS_{\min}$ | $wF_{\max}$ | $wS_{\min}$ | $wF_{\max}$ | $wS_{\min}$ |
| *Full Model* | **0.515** | **16.08** | **0.605** | **5.98** | **0.566** | **6.22** |
| Seq + Text | 0.497 | 16.85 | <u>0.594</u> | 6.14 | 0.557 | 6.45 |
| Seq + PPI | <u>0.505</u> | <u>16.18</u> | <u>0.594</u> | <u>6.09</u> | <u>0.558</u> | <u>6.33</u> |
| Seq + Structure | 0.487 | 16.94 | 0.586 | 6.25 | 0.552 | 6.43 |
| Seq Only | 0.484 | 16.99 | 0.573 | 6.39 | 0.546 | 6.48 |

Among the bi-modal configurations, the relative value of each auxiliary modality varies by biological domain. For BPO, PPI is the most informative addition, while *Sequence + Structure* performs similarly to *Sequence Only*, suggesting that structural folds are less informative than interaction networks for pathway-level annotations in this setting.

For Molecular Function, text and PPI contribute comparably: both *Sequence + Text* and *Sequence + PPI* achieve the same *wF*_max_ of 0.594, though *Sequence + PPI* attains a lower *wS*_min_ (6.09 vs 6.14). Structural data also contributes meaningfully in this aspect, raising *wF*_max_ to 0.586 from the *Sequence Only* baseline of 0.573. For Cellular Component, *Sequence + PPI* (*wF*_max_ 0.558; *wS*_min_ 6.33) marginally outperforms *Sequence + Text* (*wF*_max_ 0.557; *wS*_min_ 6.45), suggesting that interaction context is at least as informative as textual metadata for localisation tasks.

Overall, these results indicate that auxiliary modalities contribute in ontology-specific ways: PPI provides the most consistent gains across ontologies, text is competitive for MFO, and structure offers smaller but still meaningful improvements. Integrating all modalities nevertheless gives the strongest overall performance, suggesting that the modalities provide complementary rather than redundant functional evidence.

### 3.3. Comparison with State-of-the-Art Methods on CAFA3

To assess Hybrid Gated Fusion on the CAFA3 benchmark (Zhou et al., 2019), we compared it against representative methods grouped by primary input modality: sequence-based methods (DeepGOCNN (Kulmanov et al., 2018), TALE (Cao and Shen, 2021), and TEMPROT (Oliveira et al., 2023)), the structure-aware method TransFun (Boadu et al., 2023), network-based methods (DeepGraphGO (You et al., 2021) and DualNetGO (Chen and Luo, 2024)), and homology-based ensemble methods (DeepGOPlus (Kulmanov and Hoehndorf, 2020), TALE+, ATGO+, TEMPROT+, and DualNetGO+). As our data splits are consistent with prior CAFA3 evaluations, performance was compared directly using *F*_max_, the only metric consistently available across all methods and the standard basis for comparison on this benchmark.

As shown in Table 2, Hybrid Gated Fusion achieves the best reported performance in two of the three ontology aspects. In BPO, the full model (ℳ_all_) reaches an *F*_max_ of 0.601, surpassing DeepGraphGO (0.597). In CCO, it attains 0.706, improving on the ensemble-based DualNetGO+ (0.695). Notably, these results are obtained from a single model with dynamic masking, rather than from an ensemble of separately trained models.

**Table 2.**
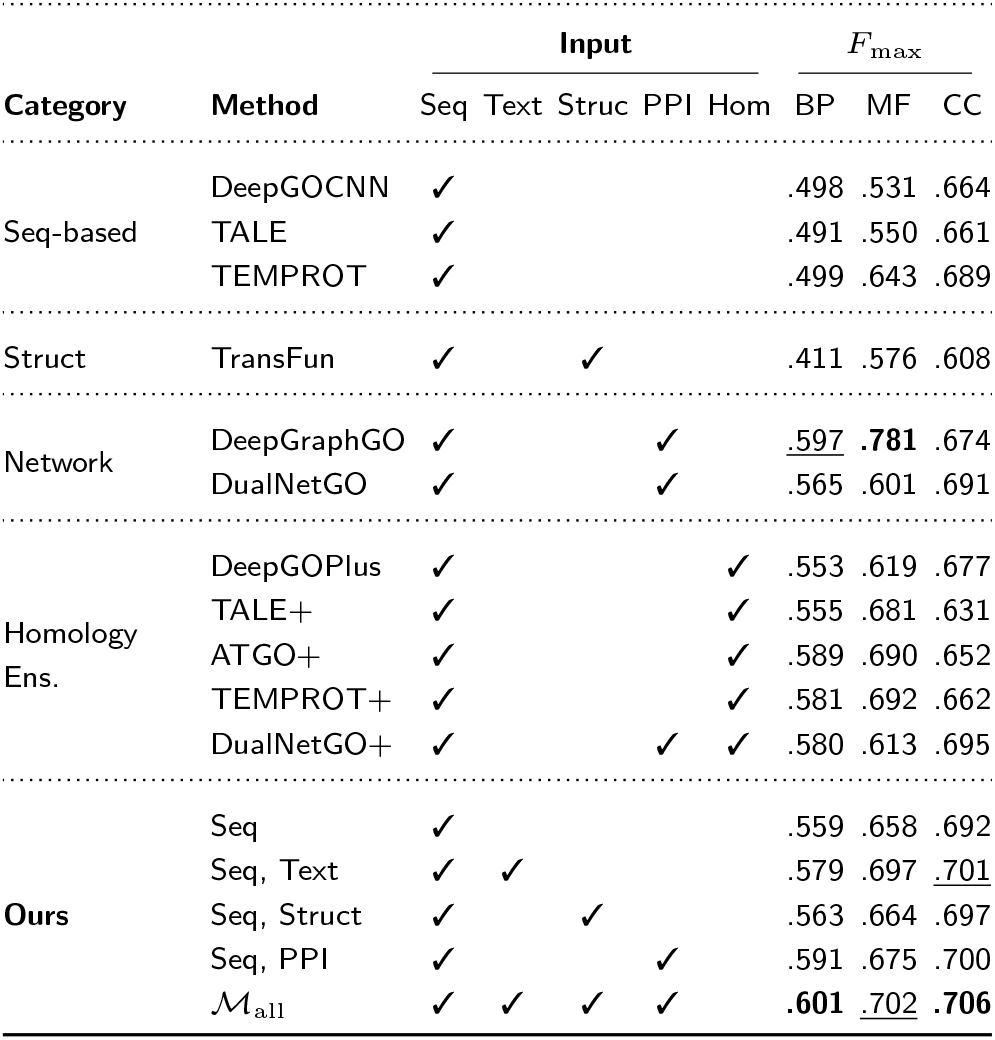
Comparative performance on CAFA3 in terms of *F*_max_. Seq, Text, Struct, PPI, and Hom denote sequence, text, structure, PPI, and homology inputs, respectively. Results for sequence-based and homologybased methods are from Oliveira et al. (2023), while those for TransFun, DeepGraphGO, and the DualNetGO variants are from Chen and Luo (2024). **Ours** denotes a single trained model with dynamic masking. Best values are shown in **bold**; second-best values are underlined.

In MFO, the model remains highly competitive, with M_all_ achieving 0.702, outperforming all sequence-based and homology-based baselines, including TEMPROT+ (0.692). DeepGraphGO retains the highest score in this aspect (0.781), likely reflecting the particular strength of PPI-derived information for molecular function prediction. Nevertheless, the Seq+Text configuration reaches 0.697, indicating that curated textual descriptions provide a strong complementary signal for function inference.

The framework is also effective under reduced-modality settings. The sequence-only configuration achieves 0.559 in BPO, outperforming dedicated sequence classifiers such as TEMPROT (0.499) and slightly exceeding DeepGOPlus (0.553), despite the latter incorporating homology information. In the structure-aware setting, the Seq+Struct configuration reaches 0.563 in BPO, substantially outperforming TransFun (0.411), suggesting that bilinear fusion with AlphaFold-derived coordinates is more effective than contact-map-based integration in this setting.

## 4. Discussion and Limitations

Hybrid Gated Fusion improves multimodal GO annotation by combining strong full-modality accuracy with substantially better robustness when evidence is incomplete. On CAFA3, a single trained model achieves state-of-the-art among the compared CAFA3 methods for BPO and CCO and remains competitive for MFO (Table 2), and is substantially more robust than early fusion when some modalities are missing at inference time (Section 3.1; Appendix G). These results suggest that the main benefit arises from learning to exploit *marginal* information across modalities, rather than simply increasing fusion capacity.

Text and PPI provide the most consistent gains over sequence across all GO aspects, with PPI contributing strongly to BPO and CCO while text remains competitive in MFO (Section 3.2). The learned gates further suggest that evidence is allocated by incremental utility: PPI is up-weighted where network context provides complementary pathway and localisation cues, whereas structure is often suppressed under full-evidence inference despite being informative in isolation (Section 3.1.2). This pattern is consistent with limited complementarity between the current structural embedding and sequence/text representations, potentially reflecting representation overlap induced by inversefolding pretraining.

Ablations support the role of the coordinated hybrid design (Section 3.1.1): auxiliary supervision preserves modality-specific discriminability, and consistency-aware aggregation aligns decision-level contributions with feature-level compatibility, reducing modality dominance and stabilising predictions without requiring separate models for different modality subsets. This is practically important because real-world annotation pipelines face uneven modality coverage.

Performance gains from structure and PPI depend on external resources (AlphaFold and STRING); dynamic masking ensures graceful degradation but cannot recover information for truly data-poor proteins. Modality missingness is also systematically biased towards well-studied proteins, which may influence learned gating and overstate benefits in low-curation regimes. Finally, we rely on frozen pretrained foundation models and evaluate primarily on CAFA3; fine-tuning these backbones with lightweight adaptation (e.g., low-rank updates) and broader stress tests would clarify robustness under wider distribution shifts.

## 5. Conclusion

We presented Hybrid Gated Fusion, a multimodal framework for GO annotation that combines compatibility-aware early gating with consistency-aware late aggregation to improve robustness under missing evidence. On CAFA3, the model achieves state-of-the-art performance for BPO and CCO in a single unified architecture and substantially improves performance in sparse-modality inference regimes. The learned gating provides an interpretable view of evidence allocation, highlighting the consistent value of curated text and interaction networks, and the regime-dependent utility of structural data. This design offers a scalable template for genomescale annotation and a modular foundation for integrating future protein representations and modalities.

## Notes

### Competing Interest Statement

The authors have declared no competing interest.

