## Appendix for "Hybrid Gated Fusion: A Multimodal Deep Learning Framework for Protein Function Annotation"

#### A Dataset Statistics

Table 3 provides a breakdown of protein counts, annotation volume, and data availability across the three GO aspects for the CAFA3 splits provided by Oliveira et al. (2023). *Avg. Terms* denotes the average number of positive annotations per protein after transitive closure propagation. *Modality Coverage* indicates the percentage of proteins possessing valid embeddings for Text, Structure, and PPI. Sequence data (ProtT5) are available for 100% of targets.

Table 3: **Dataset Statistics and Modality Coverage.** A summary of the CAFA3 benchmark splits, including dataset size, label density, and the availability of each non-sequence modality.

| | | **Size** | | **Avg.** | **Coverage (%)** | | |
|---|---|---|---|---|---|---|---|
| **Aspect** | **Split** | **#Prot** | **#Annot** | **Terms** | **Txt** | **Struc** | **PPI** |
| **BPO** | Train | 47,691 | 2,574,861 | 54.0 | 99.9 | 97.6 | 83.4 |
| | Val | 5,252 | 285,772 | 54.4 | 99.9 | 97.9 | 83.1 |
| | Test | 2,392 | 71,586 | 29.9 | 94.7 | 94.1 | 87.0 |
| **CCO** | Train | 45,309 | 779,064 | 17.2 | 99.9 | 97.8 | 88.2 |
| | Val | 4,985 | 83,840 | 16.8 | 100.0 | 98.1 | 88.0 |
| | Test | 1,265 | 16,051 | 12.7 | 91.2 | 90.6 | 90.9 |
| **MFO** | Train | 32,421 | 371,041 | 11.4 | 99.9 | 98.1 | 82.1 |
| | Val | 3,587 | 41,081 | 11.5 | 99.9 | 98.0 | 82.0 |
| | Test | 1,137 | 9,592 | 8.4 | 89.6 | 89.6 | 83.5 |

#### B Feature Extraction Implementation Details

All features are mapped to UniProt accessions to ensure consistency across splits. Preprocessing scripts were implemented in PyTorch and executed on a single NVIDIA GPU.

##### B.1 Feature Dimensions and Preprocessing

**Protein Sequence.** We extract embeddings using the `Rostlab/prot_t5_xl_half_uniref50-enc` checkpoint (Elnaggar et al., 2022).

- **Input processing:** Sequences are truncated to $L_{\max} = 1024$. Non-standard amino acids (e.g., U, Z, O, B) are normalised to 'X'.
- **Pooling:** Mask-aware mean pooling over residues produces $\mathbf{e}^{(\text{seq})} \in \mathbb{R}^{1024}$.

**Structural Features.** AlphaFold-predicted PDB files are encoded with the `esm_if1_gvp4_t16_142M_UR50` inverse-folding model (Hsu et al., 2022), based on Geometric Vector Perceptrons (GVPs) (Jing et al., 2021).

- **Input processing:** We retain backbone coordinates (N, C$\alpha$, C) only; amino-acid sequence is not provided to keep the signal geometric.
- **Pooling:** Mean pooling over per-residue outputs yields $\mathbf{e}^{(\text{struct})} \in \mathbb{R}^{512}$.

**Functional Text.** We encode UniProtKB descriptions with the Microsoft BiomedNLP PubMedBERT checkpoint (*base-uncased-abstract-fulltext*) (Gu et al., 2021).

- **Temporal decontamination:** UniProt records are continuously enriched with functional knowledge after initial deposition. Using current records for test set proteins in a temporally split benchmark would allow post-cutoff annotations to leak into the text input. To prevent this, we retrieve historical UniProt entries from the UniSave archive (Leinonen et al., 2006) for all test set proteins, selecting the latest entry version with a release date on or before 17 February 2016 (the CAFA3 training cutoff). Train and validation proteins use their current UniProt records. Historical entries are returned in UniProt flat-text format; we parse them with field-specific regular expressions to extract the same structured fields as the JSON API used for current records.
- **Field aggregation:** Eight fields are extracted and concatenated in priority order: *Protein Name*, *Function*, *Catalytic Activity*, *Pathway*, *Similarity* (family and domain membership), *Subcellular Location*, *Subunit* (complex membership), *Gene Names*. The concatenated string is truncated to 500 characters at the nearest sentence boundary.
- **Sanitisation:** Cross-references (e.g., `RHEA:`, `CHEBI:`, `GO:`), EC numbers, and evidence tags (e.g., `{ECO:...}`) are removed from all records. Historical records additionally undergo conservative punctuation normalisation to correct stitching artefacts introduced by flat-text field assembly.
- **Pooling:** We use the final `[CLS]` representation $\mathbf{e}^{(\text{text})} \in \mathbb{R}^{768}$.

**PPI Networks.** PPI embeddings are retrieved from a pre-computed STRING v12 representation learned with SPACE (Hu et al., 2025; Szklarczyk et al., 2023).

- **Mapping:** CAFA identifiers are mapped to STRING IDs via UniProt cross-references.
- **Dimension:** The interaction embedding is $\mathbf{e}^{(\text{ppi})} \in \mathbb{R}^{512}$.

### C Evaluation Metrics and Definitions

#### C.1 Information Accretion (IA)

To compute the weighted metrics, we define a weight $ia(v)$ for each Gene Ontology term $v$. $ia(v)$ corresponds to the **Information Accretion** as formally defined by Clark and Radivojac (2013). IA measures the *novel* biological information a term adds to an annotation given that its parents are already confirmed. We calculate this using the training set annotations $\mathcal{D}_{train}$:

$$ia(v) = -\log_2 \left(\Pr(v \mid \mathcal{P}(v))\right) \approx -\log_2 \left(\frac{\text{Count}(v) + 1}{\text{Count}(\mathcal{P}(v)) + 1}\right) \tag{8}$$

where:

- $\text{Count}(v)$ is the number of proteins annotated with term $v$.

- Count($\mathcal{P}(v)$) is the number of proteins annotated with **all** direct parents of $v$.
- A pseudo-count of 1 is added for Laplace smoothing.

For standard unweighted metrics, we assign $ia(v) = 1$ for all terms.

##### C.2 Metric Definitions

Using the weights defined above, Precision ($wPr$) and recall ($wRc$) are computed in a protein-centric manner. For a threshold $\tau$, let $P_i(\tau)$ and $T_i$ be the predicted and ground-truth terms for protein $i$:

$$wPr(\tau) = \frac{1}{m(\tau)} \sum_{i=1}^{m(\tau)} \frac{\sum_{f \in P_i \cap T_i} ia(f)}{\sum_{f \in P_i} ia(f)} \tag{9}$$

$$wRc(\tau) = \frac{1}{N} \sum_{i=1}^{N} \frac{\sum_{f \in P_i \cap T_i} ia(f)}{\sum_{f \in T_i} ia(f)} \tag{10}$$

where $m(\tau)$ is the number of proteins with at least one prediction. $wF_{\max}$ is the maximum harmonic mean over all thresholds:

$$wF_{\max} = \max_{\tau} \frac{2 \cdot wPr(\tau) \cdot wRc(\tau)}{wPr(\tau) + wRc(\tau)} \tag{11}$$

$wS_{\min}$ is defined as the minimum Euclidean distance between the error components (weighted Remaining Uncertainty and Misinformation):

$$wS_{\min} = \min_{\tau} \sqrt{\left(\frac{1}{N}\sum_{i=1}^{N} \sum_{f \in T_i \setminus P_i} ia(f)\right)^2 + \left(\frac{1}{N}\sum_{i=1}^{N} \sum_{f \in P_i \setminus T_i} ia(f)\right)^2} \tag{12}$$

### D Concatenation Fusion Baseline Details

To establish a standard early-fusion baseline, we implemented a Concatenation Fusion module. Unlike gated or attention-based methods which learn dynamic weights for each modality, this approach relies on a dense projection to learn static feature combinations.

**Feature Flattening.** Given the input modality features $H \in \mathbb{R}^{N \times d}$, we first flatten the modality dimension to create a single concatenated feature vector $H_{flat}$. Note that inputs $H$ are pre-masked such that $H_i = \mathbf{0}$ if the corresponding mask $M_i = 0$. The flattening operation is defined as:

$$H_{flat} = \text{Concat}(H_1, H_2, \ldots, H_N) \in \mathbb{R}^{N \cdot d}$$

**Dimensionality Reduction.** The high-dimensional concatenated vector is projected back to the common hidden space dimension $d$ using a learnable linear transformation. This layer allows the model to capture non-linear interactions between fixed positions in the modality vector (e.g., correlations between the $i$-th dimension of the sequence embedding and the $j$-th dimension of the structure embedding):

$$z = W_{proj} H_{flat} + b_{proj}$$

where $W_{proj} \in \mathbb{R}^{d \times (N \cdot d)}$ and $b_{proj} \in \mathbb{R}^d$ are learnable parameters.

### E Fusion mechanism ablation study (weighted metrics)

This section reports the full weighted evaluation for the fusion ablation described in the main text. We compare the proposed **Hybrid Gated** model against controlled variants that isolate the early pathway (*Bilinear Gated*), replace bilinear interactions with a dense projection (*Concatenation*), isolate the late pathway (*Aux Head Only*), and form an uncoordinated hybrid baseline (*Concat + Aux Head Only*). Metrics are reported for Biological Process (BPO), Molecular Function (MFO), and Cellular Component (CCO) using weighted maximum F-measure ($wF_{\max}$) and weighted minimum semantic distance ($wS_{\min}$). Unweighted results are provided in Appendix F.

Table 4: **Ablation Study Results (weighted).** Comparison of fusion configurations. **Metrics:** $wF_{\max}$ (higher is better ↑) and $wS_{\min}$ (lower is better ↓). Best results are **bold**; second-best are <u>underlined</u>.

| | | BPO | | MFO | | CCO | |
|---|---|---|---|---|---|---|---|
| **Cat.** | **Model Configuration** | $wF_{\max}$ | $wS_{\min}$ | $wF_{\max}$ | $wS_{\min}$ | $wF_{\max}$ | $wS_{\min}$ |
| **Hybrid Fusion** | Hybrid Gated (Full) | **0.515** | **16.08** | **0.605** | **5.98** | **0.566** | **6.22** |
| | Concat + Aux Late | <u>0.505</u> | 16.86 | <u>0.602</u> | 6.44 | 0.551 | 6.50 |
| Early Fusion | Bilinear Gated | 0.504 | <u>16.53</u> | 0.599 | 6.24 | <u>0.560</u> | <u>6.35</u> |
| | Concatenation | 0.500 | 16.62 | 0.586 | 6.30 | 0.553 | 6.38 |
| Late Fusion | Aux Head Only | 0.502 | 16.55 | <u>0.602</u> | <u>6.01</u> | 0.556 | <u>6.35</u> |

### F Analysis of Fusion Mechanisms (Unweighted Metrics)

In the main text, we focused on weighted metrics ($wF_{\max}$ and $wS_{\min}$) as they incorporate the hierarchical information content of Gene Ontology terms. For completeness, we report the corresponding unweighted metrics ($F_{\max}$ and $S_{\min}$) in Table 5.

Overall, the unweighted results corroborate the weighted analysis: the proposed *Hybrid Gated* model achieves the highest $F_{\max}$ across all three ontologies and the lowest $S_{\min}$ in MFO and CCO. In BPO, the *Aux Head Only* variant attains a marginally lower $S_{\min}$ (18.26 vs 18.28), but the difference is negligible. These results confirm that the overall conclusions are not sensitive to whether information-content weighting is applied.

Table 5: **Ablation Study Results (Unweighted).** Comparison of fusion configurations using standard metrics. **Metrics:** $F_{\max}$ (higher is better ↑) and $S_{\min}$ (lower is better ↓). Best results are **bold**; second-best are <u>underlined</u>.

| | | BPO | | MFO | | CCO | |
|---|---|---|---|---|---|---|---|
| **Cat.** | **Model Configuration** | $F_{\max}$ | $S_{\min}$ | $F_{\max}$ | $S_{\min}$ | $F_{\max}$ | $S_{\min}$ |
| **Hybrid Fusion** | Hybrid Gated (Proposed) | **0.601** | <u>18.28</u> | **0.702** | **4.17** | **0.706** | **6.06** |
| | Concat + Aux Late | 0.591 | 18.70 | <u>0.692</u> | 4.33 | 0.697 | 6.23 |
| Early Fusion | Bilinear Gated | <u>0.593</u> | 18.43 | 0.691 | 4.33 | 0.703 | 6.16 |
| | Concatenation | 0.589 | 18.86 | 0.683 | 4.39 | 0.697 | 6.26 |
| Late Fusion | Aux Head Only | 0.591 | **18.26** | 0.689 | <u>4.23</u> | <u>0.704</u> | **6.06** |

### G Performance on Bilinear Gated and the Hybrid Gated Fusion mode (weighted Metrics)

This section reports the full weighted evaluation for the masked-modality ablation described in the main text. We compare the **Bilinear Gated** early-fusion baseline to the full **Hybrid Gated** model under inference-time modality masking. Metrics are reported for Biological Process (BPO), Molecular Function (MFO), and Cellular Component (CCO) using weighted maximum F-measure ($wF_{\max}$) and weighted minimum semantic distance ($wS_{\min}$). Unweighted metrics are provided in Appendix I.

Table 6: **Performance comparison on the CAFA3 benchmark under modality masking.** Weighted Maximum F-measure ($wF_{\max}$) and weighted minimum semantic distance ($wS_{\min}$) are reported for BPO, MFO, and CCO. **Modality** indicates the subset available at inference. **Cov.** denotes approximate modality coverage in the dataset. Best results within each row and ontology are in **bold**.

| | | BPO | | | | MFO | | | | CCO | | | |
|---|---|---|---|---|---|---|---|---|---|---|---|---|---|
| | | Bilinear Gated | | **Hybrid Gated** | | Bilinear Gated | | **Hybrid Gated** | | Bilinear Gated | | **Hybrid Gated** | |
| **Modality** | **Cov.** | $wF_{\max}$ | $wS_{\min}$ | $wF_{\max}$ | $wS_{\min}$ | $wF_{\max}$ | $wS_{\min}$ | $wF_{\max}$ | $wS_{\min}$ | $wF_{\max}$ | $wS_{\min}$ | $wF_{\max}$ | $wS_{\min}$ |
| {Seq} | 100% | **0.458** | **17.41** | 0.452 | 17.59 | 0.537 | 7.64 | **0.566** | **6.68** | 0.532 | 6.73 | **0.539** | **6.65** |
| {Text} | ~90% | 0.399 | 19.51 | **0.447** | **17.67** | 0.474 | 8.44 | **0.539** | **6.82** | 0.492 | 7.23 | **0.510** | **6.92** |
| {Struct} | ~90% | 0.256 | 22.92 | **0.424** | **17.85** | 0.286 | 9.73 | **0.508** | **7.05** | 0.408 | 7.81 | **0.462** | **7.23** |
| {PPI} | ~85% | 0.383 | 21.02 | **0.463** | **17.92** | 0.392 | 9.89 | **0.490** | **7.27** | 0.480 | 7.06 | **0.517** | **6.60** |
| {Seq, Text} | — | 0.472 | 17.40 | **0.491** | **17.00** | 0.580 | 6.79 | **0.600** | **6.14** | 0.549 | 6.59 | **0.555** | **6.42** |
| {Seq, Struct} | — | **0.458** | **17.39** | **0.458** | 17.50 | 0.544 | 7.53 | **0.571** | **6.62** | 0.534 | 6.77 | **0.545** | **6.59** |
| {Seq, PPI} | — | 0.492 | 17.05 | **0.498** | **16.28** | 0.560 | 7.24 | **0.581** | **6.33** | 0.549 | 6.42 | **0.554** | **6.31** |
| $\mathcal{M}_{all}$ | — | 0.504 | 16.53 | **0.515** | **16.08** | 0.599 | 6.24 | **0.605** | **5.98** | 0.560 | 6.35 | **0.566** | **6.22** |

### H Performance on Bilinear Gated and the Hybrid Gated Fusion mode (Unweighted Metrics)

Table 7 presents the unweighted F-measure ($F_{\max}$) and semantic distance ($S_{\min}$) for all experimental configurations. While the main text reports weighted metrics to account for the information content (IC) of specific GO terms, the unweighted metrics are provided here for direct comparison with prior literature that may not utilise IC-weighted evaluation. Consistent with the weighted results, the **Hybrid Gated Fusion** model demonstrates superior robustness in data-sparse regimes and competitive performance in the full-modality setting.

### I Modality Analysis (Unweighted Metrics)

Table 8 presents the unweighted performance metrics ($F_{max}$ and $S_{min}$). While the general rankings largely align with the weighted results in the main text, the performance gaps are notably compressed. The Full Model achieves the highest $F_{\max}$ across all three ontologies and the lowest $S_{\min}$ in MFO and CCO; in BPO, its $S_{\min}$ (18.28) is only marginally higher than the *Sequence + PPI* baseline (18.33).

This compression suggests that unweighted metrics are less sensitive to term specificity. By contrast, the IA-weighted results in Table 1 of the main paper make the Full Model's advantage much clearer, indicating that the narrower unweighted gaps largely reflect saturation on generic, high-frequency terms rather than a limitation of the fusion strategy.

Table 7: **Unweighted performance comparison on the CAFA3 benchmark under modality masking.** Maximum F-measure ($F_{\max}$) and minimum semantic distance ($S_{\min}$) are reported for BPO, MFO, and CCO. **Modality** indicates the subset available at inference. **Cov.** denotes approximate modality coverage in the dataset. Best results within each row and ontology are in **bold**.

| | | BPO | | | | MFO | | | | CCO | | | |
|---|---|---|---|---|---|---|---|---|---|---|---|---|---|
| | | Bilinear Gated | | **Hybrid Gated** | | Bilinear Gated | | **Hybrid Gated** | | Bilinear Gated | | **Hybrid Gated** | |
| **Modality** | **Cov.** | $F_{\max}$ | $S_{\min}$ | $F_{\max}$ | $S_{\min}$ | $F_{\max}$ | $S_{\min}$ | $F_{\max}$ | $S_{\min}$ | $F_{\max}$ | $S_{\min}$ | $F_{\max}$ | $S_{\min}$ |
| {Seq} | 100% | 0.554 | **20.05** | **0.559** | 20.06 | 0.638 | 4.76 | **0.658** | **4.72** | 0.688 | 6.54 | **0.692** | **6.41** |
| {Text} | ~90% | 0.504 | 21.96 | **0.543** | **20.96** | 0.591 | 5.42 | **0.648** | **4.71** | 0.660 | 6.90 | **0.670** | **6.80** |
| {Struct} | ~90% | 0.383 | 24.27 | **0.533** | **20.93** | 0.472 | 6.37 | **0.621** | **4.98** | 0.616 | 7.80 | **0.655** | **6.85** |
| {PPI} | ~85% | 0.482 | 22.63 | **0.549** | **21.12** | 0.523 | 5.79 | **0.607** | **5.23** | 0.661 | 7.20 | **0.678** | **7.00** |
| {Seq, Text} | — | 0.571 | 19.61 | **0.579** | **19.50** | 0.676 | 4.48 | **0.697** | **4.30** | 0.697 | 6.34 | **0.701** | **6.22** |
| {Seq, Struct} | — | 0.555 | 20.02 | **0.563** | **19.87** | 0.640 | 4.75 | **0.664** | **4.65** | 0.690 | 6.54 | **0.697** | **6.33** |
| {Seq, PPI} | — | 0.581 | 18.94 | **0.591** | **18.59** | 0.658 | 4.60 | **0.675** | **4.50** | 0.693 | 6.33 | **0.700** | **6.20** |
| $\mathcal{M}_{all}$ | — | 0.593 | 18.43 | **0.601** | **18.28** | 0.691 | 4.33 | **0.702** | **4.17** | 0.703 | 6.16 | **0.706** | **6.06** |

Table 8: **Unweighted Modality Contribution Analysis.** Comparison of model performance using standard metrics. The performance gaps are much smaller here than in the weighted analysis, indicating that unweighted metrics fail to distinguish improvements in term specificity. **Metrics:** $F_{\max}$ (↑) and $S_{\min}$ (↓). Best results are **bold**; second-best are underlined.

| | BPO | | MFO | | CCO | |
|---|---|---|---|---|---|---|
| **Modalities** | $F_{\max}$ | $S_{\min}$ | $F_{\max}$ | $S_{\min}$ | $F_{\max}$ | $S_{\min}$ |
| *Full Model* | **0.601** | **18.28** | **0.702** | **4.17** | **0.706** | **6.06** |
| Seq + Text | 0.581 | 19.19 | <u>0.688</u> | <u>4.27</u> | <u>0.702</u> | 6.23 |
| Seq + PPI | <u>0.592</u> | <u>18.33</u> | <u>0.688</u> | 4.34 | 0.701 | 6.19 |
| Seq + Structure | 0.575 | 19.40 | 0.673 | 4.37 | 0.701 | <u>6.16</u> |
| Seq Only | 0.571 | 19.38 | 0.669 | 4.49 | 0.698 | 6.26 |

### J Case studies of dominant gating

This appendix contextualizes the aggregate gating analysis in the main text by examining proteins with strongly polarized gate allocations under full-modality inference. We define a modality as *dominant* for a protein–ontology pair when it receives the largest softmax-normalized gate weight and that weight exceeds 0.6. This threshold identifies genuinely concentrated allocations rather than merely the largest of four weights. When selecting the examples below, we additionally required high per-protein $F_1$ so that the biological interpretation is attached to correct predictions; prediction quality is therefore used as a sanity filter, not as evidence for dominance itself.

Table 9: **Prevalence of dominant-gate proteins under full-modality inference.** The table reports proteins with all four modalities available and the subset whose maximum gate weight exceeds 0.6. Counts in the rightmost columns are reported within that dominant-gate subset.

| **Aspect** | **All-4** | $\max \alpha_k > 0.6$ | **Seq** | **Text** | **Struct** | **PPI** |
|---|---|---|---|---|---|---|
| BPO | 1,965 | 93 (4.7%) | 11 (11.8%) | 5 (5.4%) | 0 | 77 (82.8%) |
| MFO | 839 | 57 (6.8%) | 37 (64.9%) | 0 | 0 | 20 (35.1%) |
| CCO | 1,038 | 280 (27.0%) | 73 (26.1%) | 5 (1.8%) | 0 | 202 (72.1%) |

Table 9 shows that strongly polarized gates are uncommon in BPO and MFO, but substantially more frequent in CCO. Within this tail subset, PPI accounts for most dominant cases in BPO and CCO, whereas sequence accounts for most dominant cases in MFO; structure never exceeds the 0.6 threshold in the all-modality subset. Text-dominant proteins are rare in this cohort (five in BPO and five in CCO), so we focus the qualitative discussion on the modality–ontology pairings that account for most polarized cases. The three examples below should therefore be read as illustrative proteins that align with the cohort-level tendencies in Table 9, rather than as representative behavior for the full test set. A fuller distributional characterization of per-protein gate values is left for future work.

**ZBTB5 (BPO, PPI-dominant).** ZBTB5 (Zinc finger and BTB domain-containing protein 5; *Homo sapiens*, UniProt: O15062) illustrates a PPI-dominant case for biological process prediction. UniProt and GO annotate ZBTB5 as a reviewed nuclear transcriptional regulator, with GO terms including DNA-binding transcription factor activity, RNA polymerase II-specific, DNA-binding transcription repressor activity, RNA polymerase II-specific, negative regulation of transcription by RNA polymerase II (GO:0000122), and regulation of transcription by RNA polymerase II (GO:0006357). Experimental evidence further identifies ZBTB5 as a POK-family transcription repressor (Koh et al., 2009). For BPO prediction, the model assigns a PPI gate weight of 0.645 and attains per-protein $F_1 = 0.992$ across 67 GO terms. This pattern is consistent with process-level regulatory annotations depending more strongly on interaction context and transcriptional-complex associations than on an isolated motif signal.

**ESPNL (CCO, PPI-dominant).** ESPNL (Espin-like; *Mus musculus*, UniProt: H3BLK9) highlights the strongest PPI-dominant case we identified for cellular component prediction. Because the UniProt entry is currently unreviewed, the biological support here is best read jointly from Ebrahim *et al.*, who identify espin-like as a MYO3A/MYO3B cargo in developing stereocilia and report that it is required for normal hearing (Ebrahim et al., 2016), and from the current UniProt subcellular-location annotation and GO record, which place the protein in the stereocilium (GO:0032420). For CCO prediction, the model assigns a PPI gate weight of 0.905 and attains per-protein $F_1 = 0.954$ across 32 GO terms. This makes a strong biological match for CCO: localization is determined by the stereociliary assembly context, not by sequence features alone.

**AOP3 (MFO, sequence-dominant).** AOP3 (2-oxoglutarate-dependent dioxygenase; *Arabidopsis thaliana*, UniProt: Q93VZ6) provides the sequence-dominant contrast case for molecular function prediction. UniProt classifies AOP3 within the iron/ascorbate-dependent oxidoreductase family, and its current GO annotations include 2-oxoglutarate-dependent dioxygenase activity (GO:0016706) and glucosinolate biosynthetic process. Supporting literature also links AOP3 to glucosinolate biosynthesis in *Arabidopsis* (Kliebenstein et al., 2001). For MFO prediction, the model assigns a sequence gate weight of 0.716 and attains per-protein $F_1 = 1.000$ across 6 GO terms, which is plausible for a small annotation set centered on a conserved enzymatic family. In this setting, the dominant sequence weight matches a target whose molecular function is tied to a conserved catalytic dioxygenase family.

Taken together, these examples reinforce the biological distinction suggested by the cohort-level statistics: PPI-dominant gating is most plausible when the target annotation reflects process-level context or subcellular localization, whereas sequence-dominant gating is most plausible when the target annotation reflects conserved catalytic activity. The appendix cases therefore serve mainly to illustrate how the learned gates can align with ontology-specific sources of functional evidence in a small number of strongly polarized proteins.
